# Estimations of the weather effects on brain functions using functional MRI: a cautionary note

**DOI:** 10.1101/646695

**Authors:** Xin Di, Marie Wolfer, Simone Kühn, Zhiguo Zhang, Bharat B. Biswal

**Affiliations:** Department of Biomedical Engineering, New Jersey Institute of Technology, Newark, NJ, 07102, USA; School of Life Sciences and Technology, University of Electronic Science and Technology of China, Chengdu, China; Clinical Affective Neuroimaging Laboratory (CANLAB), Otto-von-Guericke-University Magdeburg, Magdeburg, Germany; Department for Behavioral Neurology, Leibniz Institute for Neurobiology, Magdeburg, Germany; Center for Lifespan Psychology, Max Planck Institute for Human Development, Berlin, Germany; Clinic and Polyclinic for Psychiatry and Psychotherapy, University Clinic Hamburg-Eppendorf, Germany; School of Biomedical Engineering, Health Science Center, Shenzhen University, Shenzhen, China; Guangdong Provincial Key Laboratory of Biomedical Measurements and Ultrasound Imaging, Shenzhen, China

**Keywords:** daylight length, environmental effects on the brain, machine learning regression, resting-state, scanner stability, temperature, weather

## Abstract

The influences of environmental factors such as weather on the human brain are still largely unknown. A few neuroimaging studies have demonstrated seasonal effects, but were limited by their cross-sectional design or sample sizes. Most importantly, the stability of the MRI scanner hasn’t been taken into account, which may also be affected by environments. In the current study, we analyzed longitudinal resting-state functional MRI (fMRI) data from eight individuals, where the participants were scanned over months to years. We applied machine learning regression to use different resting-state parameters, including the amplitude of low-frequency fluctuations (ALFF), regional homogeneity (ReHo), and functional connectivity matrix, to predict different weather and environmental parameters. For careful control, the raw EPI and the anatomical images were also used for predictions. We first found that daylight length and air temperatures could be reliably predicted with cross-validation using the resting-state parameters. However, similar prediction accuracies could also be achieved by using one frame of EPI image, and even higher accuracies could be achieved by using segmented or raw anatomical images. Finally, the signals outside of the brain in the anatomical images and signals in phantom scans could also achieve higher prediction accuracies, suggesting that the predictability may be due to the baseline signals of the MRI scanner. After all, we did not identify detectable influences of weather on brain functions other than the influences on the baseline signals of MRI scanners. The results highlight the difficulty of studying long-term effects using MRI.

## 1. Introduction

Daily environmental factors such as weather and seasonality affect mood and cognitive functions (Cedeñ o Laurent et al., 2018; Denissen et al., 2008; IJzerman et al., 2018; Keller et al., 2005; Lim et al., 2018), and may lead to pathological affective disorder (Elseoud et al., 2014; Kurlansik and Ibay, 2012). The effects on individuals may be small, but the collective effects may lead to broader impacts, e.g. on stock markets (Hirshleifer and Shumway, 2003; Saunders, 1993). To better understand the effects of weather and seasonality on mood or cognition, it is critical to study their effects on brain functions. A few human neuroimaging studies have explored this association. Seasonal effects on brain functions as measured by functional MRI (fMRI) have been observed both in resting-state (Choe et al., 2015) and when performing cognitive tasks (Meyer et al., 2016). Some neural transmitter activity in the striatum also showed seasonal effects, i.e. serotonin transmitter binding as measured by ^11^C–labeled 3-amino-4-(2-dimethylaminomethyl-phenylsulfanyl)-benzonitrile ([^11^C]DASB) positron emission tomography (PET) (Kalbitzer et al., 2010; Mc Mahon et al., 2016; Praschak-Rieder et al., 2008) and dopamine synthesis as measured by ^18^F-DOPA PET (Eisenberg et al., 2010; Kaasinen et al., 2012). A study even reported seasonal changes of hippocampal volumes in human subjects (Miller et al., 2015).

There are several limitations in these neuroimaging studies. First, most of these studies are cross-sectional, which is limited by the large individual differences in brain functions (Gordon et al., 2017). In addition, most of the studies examined roughly defined seasonal effects or yearly periodical effects. But the exact phase of the seasonal variations may be different from the four seasons. Sometimes the yearly effects showed different phases (Meyer et al., 2016), suggesting more complicated relationships of environmental factors on brain functions. Therefore, it is critical to examine which environmental parameters, such as weather, have more contributions to the seasonal effects. Among different environmental parameters, daylight length and temperature represent the significant environmental differences in seasonal fluctuations. Gillihan et al have explored the weather effects on brain functions using a small cross-sectional sample (Gillihan et al., 2011). They identified a weather index related to mood and showed that the weather index was correlated with resting-state cerebral blood flow as measured by arterial spin labeling (ASL) perfusion fMRI mainly in the insula. But more systematic examinations of weather effects on brain functions have not been performed. Lastly, the most commonly used neuroimaging method is fMRI based on blood-oxygen-level dependent (BOLD) signals (Ogawa et al., 1990), where the interpretation of the results should consider neuronal level, neurophysiological level, and the underlying physical level of the scanner. Specifically, if some effects on fMRI signals were observed, they may be due to the changes in neuronal activity, which is favorable to psychologists and psychiatrists. But the effects may also due to the changes in neurovascular coupling (Di et al., 2013; Yuan et al., 2013), in brain structures, or even the stability of the MRI scanner. Therefore, when examining the weather effects on brain functions, alternative factors need to be considered and carefully controlled.

The purpose of the current study is to estimate to what extent resting-state brain functions were affected by the weather. We analyzed longitudinal resting-state fMRI data from eight individuals from three datasets, where the individuals were scanned over periods of months to years (Choe et al., 2015; Filevich et al., 2017; Poldrack et al., 2015). One challenge for estimating weather effects is that the effects may be small. Therefore, we applied a machine learning regression approach to evaluate the effects. Because multiple brain regions have been implicated in seasonal effects, e.g. basal ganglia (Kalbitzer et al., 2010; Mc Mahon et al., 2016; Praschak-Rieder et al., 2008), insula (Gillihan et al., 2011), and hippocampus (Miller et al., 2015), small regional effects may be aggregated into detectable effects using machine learning technique. We performed a within-subject prediction analysis at the single-subject level. We asked what weather parameters have the most effects on resting-state brain functions, which can be represented as high prediction accuracies in predictions of these parameters. In order to rule out possible confounding effects that might give rise to prediction, we also performed several control prediction analyses. First, we analyzed anatomical MRI images to check whether the observed prediction could be attributed to anatomical variations. Second, we checked images from phantom data to examine whether the prediction could be attributed to the stability of the MRI scanner.

## 2. Materials and methods

### 2.1. MRI datasets

Several multi-session resting-state fMRI datasets were pooled together, where the subjects were scanned over periods of months to years. The first subject was derived from the Kirby sample (Choe et al., 2015), where the single subject was scanned for 156 sessions over three and half years. The second subject was from the Myconnectome sample (Poldrack et al., 2015), where the subject was scanned 90 times over one and half years. The remaining six subjects were from the Day2day sample (Filevich et al., 2017), where the subjects were scanned over a similar span of about half a year. The detailed subject and scan information is listed in Table 1.

**Table 1.** Subject and MRI scan information.

|  | Dataset | Sex | Age | # of sessions | First scan | Last scan | # of volumes | TR (s) | Voxel size (mm<sup>3</sup>) |
| --- | --- | --- | --- | --- | --- | --- | --- | --- | --- |
| 1 | Kirby | M | 40 | 156 | 2009/12/07 | 2013/06/20 | 198 | 2 | 3 x 3 x 4 |
| 2 | MyConnectome | M | 45 | 83 | 2012/10/23 | 2014/03/11 | 500 | 1.16 | 2.4 x 2.4 x 2.4 |
| 3 | Day2day | F | 23 | 50 | 2013/07/03 | 2013/12/18 | 148 | 2 | 3 x 3 x 3.6 |
| 4 | Day2day | F | 31 | 48 | 2013/07/03 | 2014/01/08 | 148 | 2 | 3 x 3 x 3.6 |
| 5 | Day2day | F | 29 | 45 | 2013/07/03 | 2014/01/27 | 148 | 2 | 3 x 3 x 3.6 |
| 6 | Day2day | F | 24 | 46 | 2013/07/02 | 2013/12/19 | 148 | 2 | 3 x 3 x 3.6 |
| 7 | Day2day | M | 30 | 39 | 2013/07/09 | 2014/02/12 | 148 | 2 | 3 x 3 x 3.6 |
| 8 | Day2day | F | 29 | 48 | 2013/07/03 | 2014/02/20 | 148 | 2 | 3 x 3 x 3.6 |

The number of sessions represents the effective numbers after dropout due to missing data or large head motions. The numbers of volumes represent the numbers used in the analysis after removing the first several volumes.

The MRI data from the Kirby sample were scanned using a 3T Philips Achieva scanner. The data from the Myconnectome sample were scanned using a 3T Siemens Skyra scanner using a 32-channel head coil. And the data from the Day2day project were scanned using a 3T Siemens Magnetom Trio scanner using a 12-channel head coil. For each subject, resting-state fMRI data with multiple sessions were acquired. Within a subject, the resting-state fMRI were scanned using the same imaging parameters, but the parameters varied between different sites. Some essential resting-state fMRI parameters are listed in Table 1. For more details, we refer the readers to the original articles.

High-resolution anatomical MRI images were available for only a few sessions in the Kirby and Myconnectome datasets. An MRI image of one session was used to register all the functional images to standard Montreal Neurological Institute (MNI) space. For the Day2day dataset, structural MRI images were available for all the sessions. Only the structural MRI image of the last session of each subject was used to aid preprocessing of the fMRI images. All the structural images of the Day2day project were also used in the control prediction analysis.

Lastly, we obtained MRI scanner quality assurance agar phantom data from the Day2day site. The images were scanned between June 2013 and February 2014 on a weekly basis (37 sessions in total). One session’s data were dropped because of extreme variations in the images. The data were acquired using a gradient echo (GRE) sequence with the same coil as the one used for the acquisition of the human data. Two images were acquired for each session. The parameters include: TR = 2000 ms; TE = 30 ms; FOV = 22 cm; matrix = 64 × 64; slice number = 28; slice thickness = 4 mm (1 mm gap).

### 2.2. Environmental data

The MRI data were acquired from three different cities in two continents, Baltimore USA (Kirby), Austin USA (Myconnectome), and Berlin Germany (Day2day), which reflect different types of climates. The latitudes of these four cities are approximately 39 °N, 30 °N, and 52 °N, respectively. The weather data for the two US cities were downloaded from (US) National Centers for Environmental Information website (https://www.ncdc.noaa.gov/cdo-web/). The Local Climatological Data from Maryland Science Center Station and Austin Camp Mabry Station were used to represent the weather for the Kirby and Myconnectome datasets, respectively. We used the following measures, maximum and minimum temperatures (Temp_max_ and Temp_min_), air pressure (Press), wind speed (Wind), humidity (Hum), and precipitation (Prcp). For those with missing data, we also checked Daily Summaries data from the NOAA website. The weather data for the Day2day dataset were collected by the German researchers. Daily sunshine hours were not used, because they were not available for the other datasets.

We also included daylight length (Dalgt) in the current analysis. It was already available in the NOAA Local Climatological Data. For the Day2day data, we calculated the daylight length in Berlin according to its geographic location through the website of the Astronomical Applications Department of the U.S. Naval Observatory computes (http://aa.usno.navy.mil/data/docs/Dur_OneYear.php). For the Day2day dataset, there are three additional parameters that reflect local environmental variations, i.e. scanner room temperature (Temp_rm_), humidity (Hum_rm_), and scanner Helium level (He). These three parameters were also used in the prediction analysis when using the Day2day data.

### 2.3. MRI data processing

#### 2.3.1. Resting-state fMRI Preprocessing

Data processing and statistical analysis were performed using MATLAB (R2017b). SPM12 (http://www.fil.ion.ucl.ac.uk/spm/; RRID:SCR_007037) was used for fMRI data preprocessing. The first 2, 18, and 2 functional images for each session were discarded for the Kirby, Myconnectome, and Day2day datasets, respectively, remaining 198, 500, and 148 images for each session. For each subject, all the functional images were realigned to the first session. All the prediction analysis was performed in the native space of each subject. The anatomical images were coregistered to the mean functional image, and then segmented into gray matter (GM), white matter (WM), cerebrospinal fluid (CSF), and other tissues. For each subject, an intracranial volume mask was defined, and the grand mean (4-dimensional average) of the functional images was calculated for each session. For each session, the functional images were divided by the grand mean and multiplied by 100. At each voxel, Friston’s 24 head motion model (Friston et al., 1996), the first five principal components from WM signals and the first five principal components from CSF signals were regressed out, and then band-pass filtering was applied between 0.01 and 0.1 Hz. The images were not spatially smoothed, because there was no voxel-wise univariate analysis involved.

The preprocessing steps were chosen to minimize potential artifacts due to physiological noises and head motion. This may be an over-conservative choice that may compromise too many degrees of freedom of the fMRI time series (Bright et al., 2017). We also tried to reduce the number of regressors during the linear regression step. Specifically, we obtained the first two principal components of Friston’s 24 head motion variables. The regression then included the first two components of the head motion model, the first component of WM signals, and the first component of the CSF signals (2 + 1 + 1 regressors compared with 24 + 5 + 5 regressors from the main analysis). The results using the reduced regression were reported in the supplementary materials.

#### 2.3.2. ALFF, ReHo, and connectivity matrices

We calculated three resting-state parameters to represent resting-state brain functions, i.e. amplitude of low-frequency fluctuation (ALFF) (Zang et al., 2007) and regional homogeneity (ReHo) (Zang et al., 2004) to represent regional properties, and connectivity matrix to represent inter-regional connectivity property. ALFF and ReHo were calculated using the REST toolbox (REST: a toolkit for resting-state fMRI, RRID:SCR_009641) (Song et al., 2011). Essentially, ALFF calculated the power of the time series signals between 0.01 to 0.08 Hz at every voxel, resulting in an ALFF map for each session. ReHo calculated the correlations of the current voxel with the 26 neighboring voxels, which also resulted in a ReHo map for each session. The ALFF and ReHo values for each session within the subject’s GM mask were converted to a vector for further analysis. The subject-specific GM masks were defined as GM intensity greater than 0.5 based on the segmentation of the subject’s anatomical image. Because the GM masks were defined in the native spaces and the fMRI resolution varies across datasets, the number of within mask voxels also varied (from 20,780 to 55,368).

Correlation matrices were calculated among 164 regions of interest (Di and Biswal, 2019; Dosenbach et al., 2010). Spherical ROIs were first defined in MNI space with a radius of 8 mm, and then transformed into the native space for each subject. There were in total 13,366 connectivity values (164 x (164 - 1) / 2), which were converted to a vector for the prediction analysis. The correlation values were transformed into Fishers’ z scores.

#### 2.3.3. Head motion and other potential confounding variables

To minimize the confounding of head motion in the prediction analysis, we first removed sessions with large head motions. We calculated frame-wise displacement in translation and rotation directions (Di and Biswal, 2015). A session’s data with maximum frame-wise displacement greater than 1 mm or 1° were discarded. No sessions were removed in the Kirby data, and seven sessions (7.8%) were removed for the Myconnectome data. In the Day2day dataset, at most two sessions were removed for each subject. Secondly, we regressed out 24 motion variables using Friston’s head motion model, which has been shown to be effective to minimize the effects of head motion on resting-state measures (Yan et al., 2013). Lastly, mean frame-wise displacement of both directions were regressed out from a predicted environmental variable before it was entered into the prediction analysis.

#### 2.3.4. Global signal

The resting-state fMRI data have been scaled by the grand mean (4-D average) of each session to account for the baseline signal variations across sessions. However, a recent study has reported an association between global signal fluctuations and time of day (Orban et al., 2020). We, therefore, examined whether the global signal fluctuations were associated with the environmental factors, and whether accounting for the global signal fluctuations could affect the predictions of these environmental factors. We calculated the averaged ALFF value in the intracranial mask to reflect the global signal fluctuations. The global signal fluctuations were correlated with daylight length for each subject. Next, we also calculated mean ALFF (mALFF) by dividing an ALFF map by the global mean. Prediction analysis was also performed by using the mALFF maps.

#### 2.3.5. Structural MRI processing

For the Day2day dataset, the MPRAGE anatomical MRI images were available for all the sessions. Therefore, we used the anatomical images as a control condition for weather prediction. The analysis was also performed in a subject’s native space. The anatomical images from all the sessions of a subject were realigned and resliced to the image of the first session. Then each session’s image was segmented separately, and the segmented tissue probability maps of GM, WM, and CSF were obtained. We defined GM, WM, and CSF masks as an averaged probability greater than 0.5 for respective tissue types. GM, WM, and CSF probability in their masks were extracted, respectively, to be used in the prediction analysis.

We also defined an air mask to study the baseline MRI signals, which was located outside the brain (Makedonov et al., 2013). The mask was placed at the lower left front side of the head to avoid potential objects in the area, and was consisted of 21 × 41 × 41 voxels.

#### 2.3.6. Phantom image processing

For each session, the two images were realigned, and an averaged image was calculated. Because the phantom was imaged in a similar location, no cross-session registration was performed. We first calculated correlations between daylight length and image values in every voxel, resulting in a correlation image. Next, a cubic mask in the center of the image was defined. The signals within the mask were extracted for the prediction analysis.

### 2.4. Prediction analysis

#### 2.4.1. Prediction analysis scheme

The goal of the analysis is to estimate the prediction values of resting-state parameters on different weather or meteorological parameters. The analysis was performed for each of the resting –state parameters to predict each of the seven weather parameters. And we asked which weather parameters could be better predicted by which resting-state parameters. The prediction analysis was all done in a within-subject manner. Cross-validation was used to evaluate the prediction accuracies.

In addition to use these resting-state parameters, we also performed a series of control analyses to use other potential confounding parameters to predict the environmental parameters. First, we used the first fMRI image of each session after realignment to perform prediction analysis. Although the single image still reflects BOLD effects, brain structures may contribute more variations. Secondly, to future rule out the structural contribution, we used segmented tissue probabilities of GM, WM, and CSF from their respective tissue masks to perform prediction analysis. Thirdly, we also extracted raw image values from the MPRAGE images in the three tissue masks to serve as another control condition. In addition, a cuboid mask was defined for each subject, which was located outside the brain. The raw image values from the MPRAGE images from the air mask were used to control for baseline MRI signals. Finally, since all of the above-mentioned analyses indicated prediction values to predict environmental parameters, especially daylight length, we further analyzed the quality assurance phantom data, and used the signals in the agar phantom area to perform prediction analysis to predict daylight length.

#### 2.4.2. Machine learning regression analysis

We used a linear machine learning regression model to perform prediction analysis. The general form of the prediction model is a linear regression model as the following:

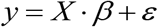

where *y* is a *n x 1* vector of a predicted weather parameter, *X* is a *n x m* matrix of a resting-state parameter, *β* is the model parameters, and *ε* is the residual. N represents the number of observations, which in the current analysis was the number of sessions for a particular subject. M represents the number of prediction variables, which could be the number of voxels in the ALFF or ReHo maps (see Table 1) or the number of connections (13,366) in the connectivity matrices. Here, m is much larger than n. Therefore, we used ridge regression to estimate the *β* parameters. Briefly speaking, instead of trying to achieve the goal of minimizing the sum of square means of the model prediction :

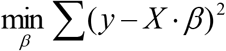

Ridge regression adds one more regularized term:

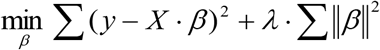

where λ represents the regularization parameter. The regularization term can constrain the sizes of beta values, thus preventing overfitting of the model. In the current analysis, we used the MATLAB function fitrlinear to perform the prediction analysis. There are other methods available, such as LASSO and elastic net, but a recent study suggested that ridge regression and elastic net can yield similar prediction accuracies while LASSO might perform worse in the scenario that the number of observations is much smaller than the number of features (Cui and Gong, 2018).

There are three steps in the prediction analysis, 1) tuning the regularization parameter λ to find the optimal λ (λ tuning), 2) training the model using the training dataset and the optimal λ to obtain a prediction model β (model training), and 3) estimating prediction accuracy by calculating correlations between predicted and actual values in a separate testing sample (cross-validation). Cross-validation was used to make sure that the estimated prediction accuracies were independent of the training data.

Because of the limited number of data in one fold (13 observations in the least case), 3-fold cross-validation was adopted. We used a nested tuning strategy to optimize the parameter λ (Cui and Gong, 2018). Specifically, we first held out one-third of the data as an independent testing dataset, and used the remaining two-thirds of the data as a training and parameter tuning dataset. The data were first sorted according to the tested weather parameter, and the three folds were defined as the 1 ^st^, 4^th^, 7^th^, …, 2^nd^, 5^th^, 8^th^, …, and 3^rd^, 6^th^, 9^th^, … sessions of the data, respectively. Within the two-thirds training and parameter tuning dataset, we first performed a nested loop of 3-fold analysis. Specifically, one-third of the data were holden out, and the remaining two-thirds of data were used to train the regression model using a set of *λ* values, from 10^−5^ to 10^−1^ in the logarithmic scale with a total of 15 values. The inner loop testing data was used to test the accuracy of the prediction by calculating the correlation between predicted and actual weather parameter values. This procedure was performed three times for the three folds, and the mean accuracies were calculated for each of the λ values. The λ value with the highest mean accuracy was used for the outer layer training data to train the model. The model was then applied to the outer layer testing data to estimate prediction accuracies. The three accuracy values from the 3 folds were averaged to represent an estimate of accuracy for a subject.

The prediction accuracies of different imaging parameters and environmental parameters were visualized by using notBoxPlot (https://github.com/raacampbell/notBoxPlot). The plot shows not only the individuals’ prediction accuracies, but also the mean, standard deviation, and 95% confidence interval of the accuracies across the subjects. False discovery rate (FDR) was used to correct for multiple comparisons of the different parameters.

## 3. Results

### 3.1. Predictions using the resting-state images

We first performed predictions on different environmental parameters using the ALFF maps, ReHo maps, connectivity matrices, as well as using a single frame of EPI images as a control condition (Figure 1). In general, daylight length (Dalgt) and maximum and minimum environmental temperatures (Temp_max_ and Temp_min_) had higher prediction accuracies, with daylight length usually having the highest prediction accuracies. The other environmental parameters had very low prediction accuracies. In terms of the resting-state parameters, the ALFF map usually had the highest prediction values. The average prediction accuracy of daylight length using ALFF was 0.38. Surprisingly, however, using a single frame of EPI images could achieve comparable and even higher prediction accuracies than any resting-state parameters. The averaged prediction accuracy of daylight length using the raw EPI images was 0.42. When using the resting-state parameters calculated from the reduced preprocessing method to perform the predictions, the prediction accuracies slightly increased (Supplementary Figure S1). However, they were still smaller than those using the raw EPI images.

**Figure 1.**
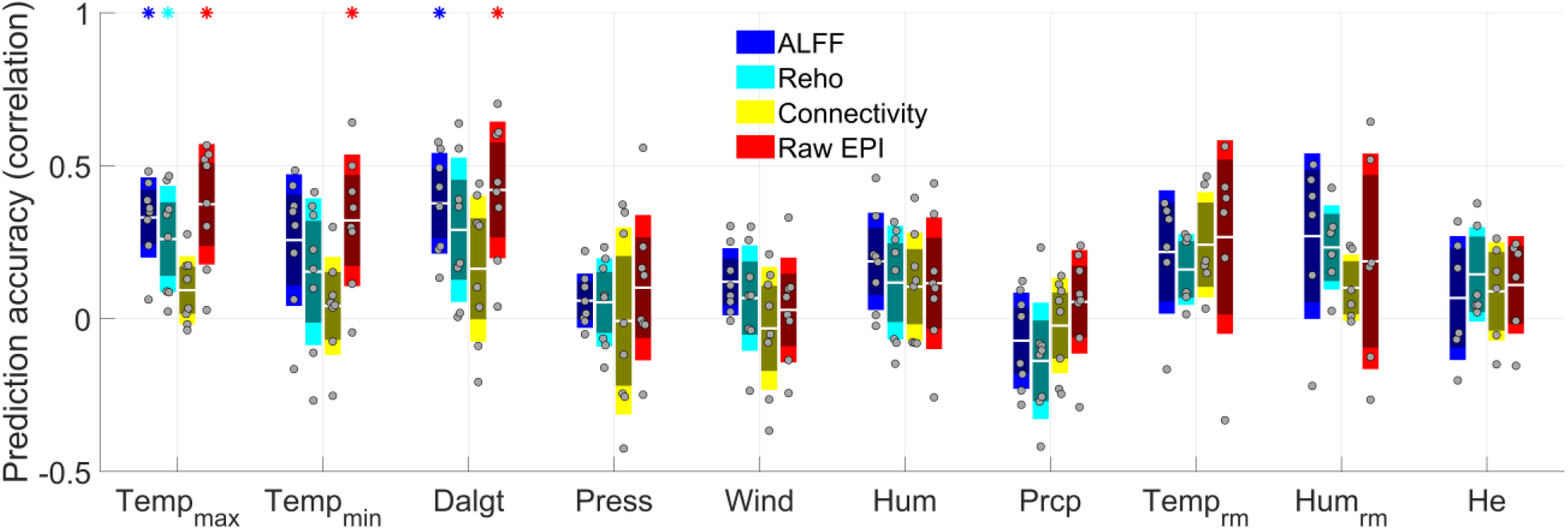
Prediction accuracies (correlations) of the amplitude of low-frequency fluctuations (ALFF) maps, regional homogeneity (ReHo) maps, connectivity matrices, and raw echo-planar imaging (EPI) maps on different environmental parameters. Each dot represents one subject’s mean prediction accuracy. The center white lines, inner dark bars, and outer light bars represent the mean, 95% confidence interval, and standard deviation, respectively. The asterisks on the top represent statistical significance at p < 0.05 after false discovery rate (FDR) correction for all the 40 predictions.

It remains a question that whether the weather predictions using the resting-state parameters and single EPI images are based on similar or different information. Since the daylight length had the highest prediction accuracies, we focused on its prediction. We combined ALFF with EPI and ReHo with EPI to predict daylight length to check whether combining the two modalities can boost the prediction accuracies. Unfortunately, combing two modalities yielded very similar prediction accuracies as those using single EPI images or ALFF images (Figure 2). Therefore, ALFF and ReHo did not convey more information than a single EPI image to predict daylight length.

**Figure 2.**
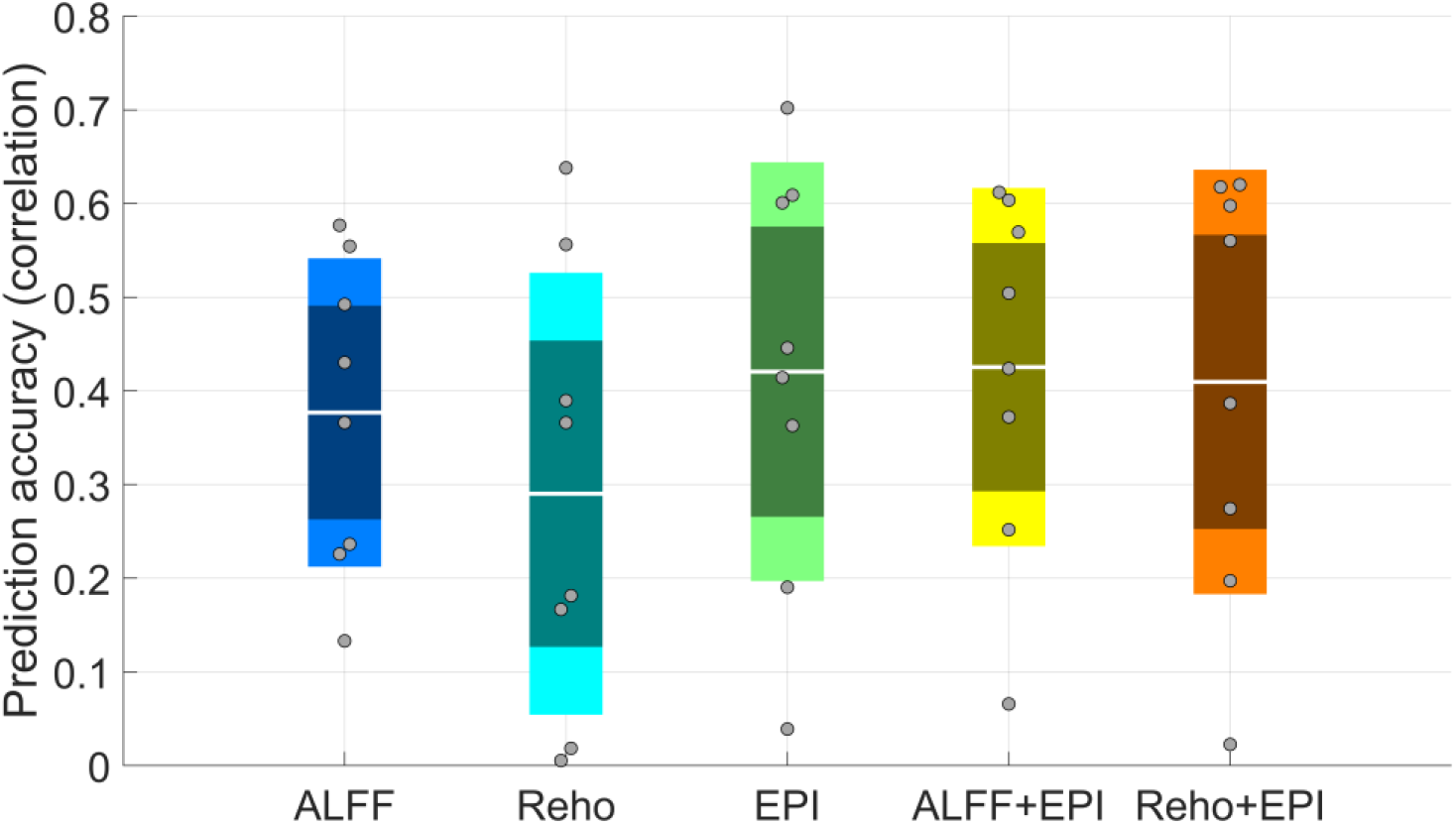
Prediction accuracies to daylight length using the amplitude of low-frequency fluctuations (ALFF), regional homogeneity (ReHo), raw echo-planar imaging (EPI) maps, and their combinations. Each dot represents one subject’s mean prediction accuracy. The center white lines, inner dark bars, and outer light bars represent the mean, 95% confidence interval, and standard deviation, respectively.

Next, we examined whether the global signal fluctuations of the resting-state data were correlated with daylight lengths, and whether the global signal fluctuations contribute to the predictions of the environmental factors. The correlations between global mean ALFF and daylight lengths did not show a consistent pattern across subjects (Figure 3A). We also used the global mean scaled ALFF maps, i.e. mALFF, to predict different environmental variables, and they yielded similar prediction patterns as what using the raw ALFF maps (Figure 3B).

**Figure 3.**
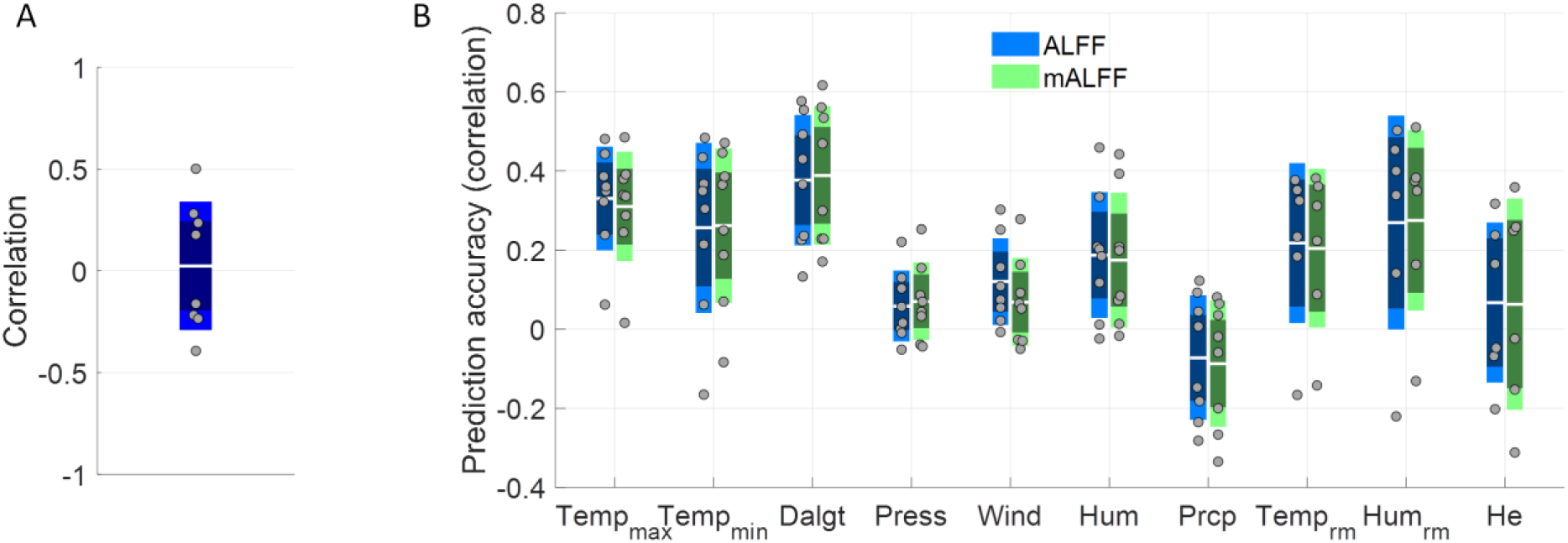
A, correlations between the global amplitude of low-frequency fluctuations (ALFF) and daylight lengths. B, prediction accuracies (correlations) of the environmental parameters using raw ALFF and mean ALFF (mALFF). Each dot represents one subject’s mean prediction accuracy. The center line, inner dark bar, and outer light bar represent the mean, 95% confidence interval, and standard deviation, respectively.

### 3.2. Predictions using the anatomical images

If a single volume of EPI image can predict weather parameters like daylight length, then the question becomes whether the prediction is due to brain functional activity, structural information, or other factors. We, therefore, performed similar prediction analyses using the anatomical images, which are available in the six subjects in the Day2day dataset. We first performed predictions using the segmented GM, WM, or CSF density images within their respective tissue masks (Figure 4). The results showed very similar prediction patterns for different environmental parameters as what using the resting-state parameters. That is, the daylight length and environmental temperatures had the highest prediction accuracies. The prediction accuracies using all the three tissue probability maps were above 0.5, which were higher than using any of the resting-state parameters. However, what was more interesting was that even higher prediction accuracies could be achieved using the raw MRI signals in these tissue masks. The prediction accuracies were higher than 0.6 when using raw MRI signals in the GM and CSF masks. Finally, we defined a cuboid mask outside the brain (see Figure 5A as an example), and used the raw MRI signals in the mask to perform prediction analysis. Surprisingly, the analysis also showed a similar pattern of prediction accuracies. The prediction accuracy on daylight length using the air mask was 0.47, which was lower than using all the other anatomical parameters but still higher than using any of the resting-state parameters.

**Figure 4.**
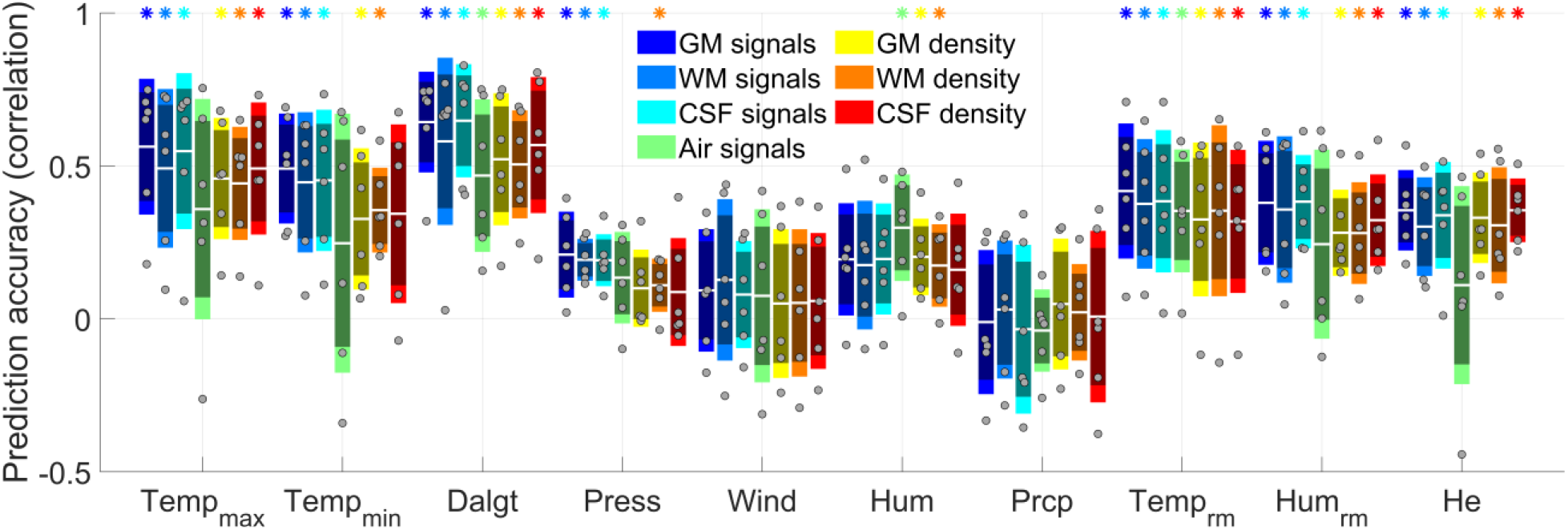
Prediction accuracies (correlations) of raw MRI signals and segmented densities in gray matter (GM), white matter (WM), cerebrospinal fluid (CSF), and air masks on different environmental parameters. Each dot represents one subject’s mean prediction accuracy. The center white lines, inner dark bars, and outer light bars represent the mean, 95% confidence interval, and standard deviation, respectively. The asterisks on the top represent statistical significance at p < 0.05 after false discovery rate (FDR) correction for all the 70 predictions.

**Figure 5.**
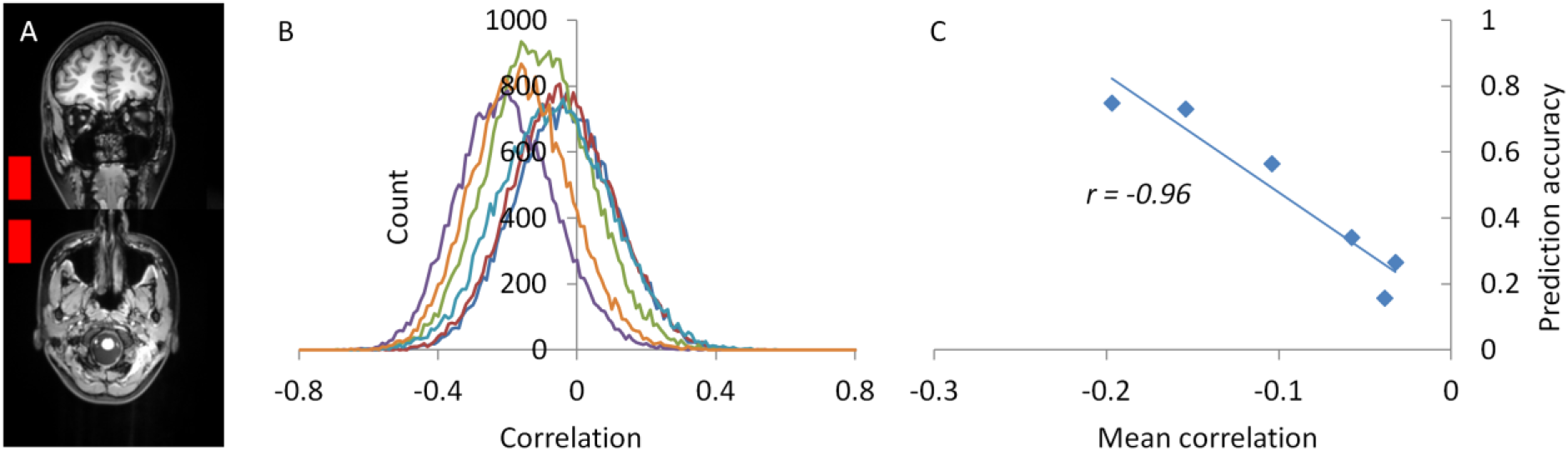
A) An example of the air mask from one subject overlaid to the subject’s anatomical image. B) Histograms of the correlations between the MRI signals and daylig ht length of all the voxels in the air mask. Each line represents one subject. C) There is an extremely high negative correlation between the mean correlations in the air mask and the prediction accuracies of using the air mask voxels to predict daylight length.

To further explore the baseline MRI signals conveyed in the air mask, we calculated correlations between the MRI signals and daylight length in all the voxels in the air mask for the six subjects (Figure 5B). There were small global effects of correlations between the MRI signals and daylight lengths. Moreover, the global effects of correlations were strongly correlated with the prediction accuracies across subjects (Figure 5C), indicating that the global correlation is the driving information that gave rise to the prediction accuracy.

### 3.3. Control analysis using the phantom images

To further confirm the baseline signal changes, we analyzed the weekly quality control phantom data around the same period of the Day2day project. We first calculated voxel-wise correlations between the MRI signal and daylight lengths (Figure 6A). It clearly showed that in the phantom region, there were high negative correlations. We defined a cubic mask in the center of the image, and the distribution of correlations of all the voxels in the mask is plotted in Figure 6B. The mean and median correlation in the cubic mask was *-0*.*78* and *-0*.*79*, respectively. We also performed similar predictions of the daylight length by using the MRI signals in the mask, and the cross-validated mean accuracy was *0*.*70*. Lastly, we calculated the mean and spatial coefficient of variation of the MRI signals in the mask, and plotted them against scan sessions (Figure 6C and 6D). The mean MRI signals showed a strong negative correlation with daylight lengths (*r = -0*.*82, p < 0*.*001*). However, the spatial coefficient of variation showed only a marginally significant correlation (*r = -0*.*34, p = 0*.*04*). The large negative correlation of daylight length with the mean MRI signals and reduced correlation with the spatial coefficient of variation were also confirmed by using the air mask signals of the MPRAGE images (Supplementary Figure S2).

**Figure 6.**
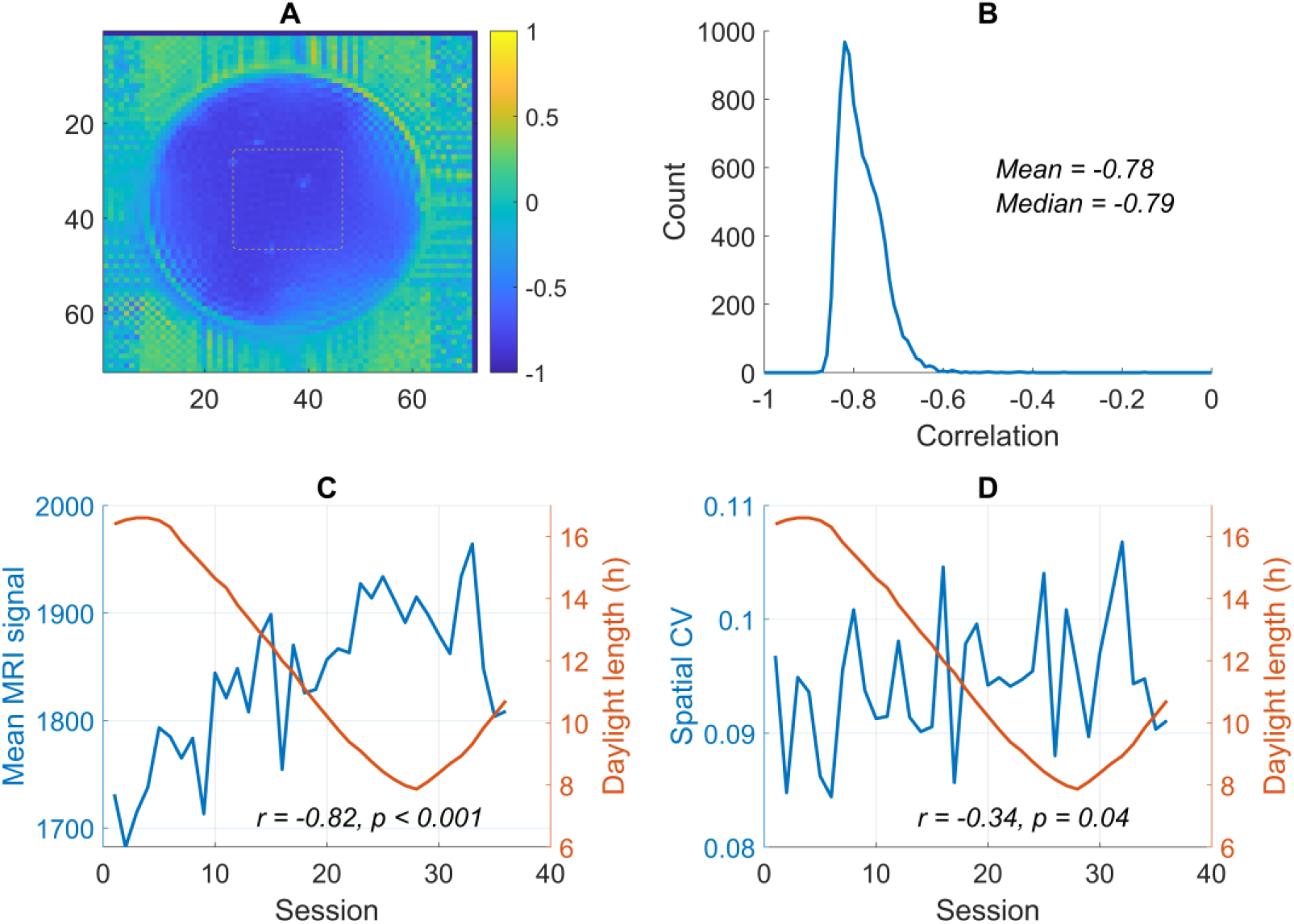
A) Voxel-wise correlation maps between MRI image values and daylight length. The dash-line contour indicates the location of the cubic mask. B) Histogram of the voxel-wise correlations in the cubic mask. C) and D) The averaged signals and spatial coefficient of variation (CV) in the cubic mask and the daylight length against scan sessions.

## 4. Discussion

By applying machine learning regression to single-subject longitudinal fMRI data that were scanned over months to years, we demonstrated that we can predict environmental parameters, especially daylight length and air temperature, by using resting-state fMRI parameters. However, a series of controlled analyses showed that using a single EPI image, the segmented tissue density, and the raw MRI signals from the anatomical images in different tissue masks and even in a mask outside the brain could all predict the environmental parameters. The resting-state parameters did not add prediction values to single-volume EPI images. These results indicated the prediction of environmental parameters, especially daylight length, cannot be explained as the weather effects on brain functions. Rather, the prediction may reflect MRI scanner baseline signal variations that were affected by the environmental parameters. The analysis of the quality control phantom images supported our speculation.

Among all the environmental parameters analyzed the daylight length and air temperature had the highest prediction accuracies. It is not surprising because daylight length and air temperature are highly correlated. Daylight length has the highest prediction accuracy probably because it is a physical quantity that does not have measurement errors, which is in contrast to air temperature. It’s noteworthy that although the MRI scanner room temperature and humidity could not be predicted by the functional parameters, they could be reliably predicted by the anatomical MRI parameters (Figure 4). However, their prediction accuracies were smaller than those of daylight length and air temperature. It indicates that the environmental effects on MRI signals are not directly caused by local temperature, but some other local factors. A study has shown that the gaseous oxygen level in the magnet field can influence the MRI signals (Bates et al., 1995). The oxygen level in the scanner room may fluctuate across seasons due to different ventilation conditions, which may contribute to the MRI baseline signal shifts. In addition, the cooling systems of the scanner may be affected by either electricity supply stability or cooling water temperature. Given that the MRI is such a sophisticated machine, there may be other factors that mediate the association between daylight length and scanner stability.

The current results highlighted the difficulty to study long-term effects such as weather on brain structures and functions using MRI. Consistent with two previous fMRI studies (Choe et al., 2015; Meyer et al., 2016), we did find weather effects on fMRI measures. But we demonstrated that the weather effects are likely due to the variations of MRI scanner baseline. It is reasonable to speculate that MRI scanner stability might contribute to the reported seasonal effects (Choe et al., 2015; Meyer et al., 2016). We also showed that tissue probability measures of GM volumes may also be affected by the scanner stability, so brain volumetric measures may also be affected by the scanner stability (Miller et al., 2015). Careful examinations of the effects of MRI scanner baseline signals are needed to confirm these reported findings.

In the current analysis, a few steps have been used to correct the MRI baseline signals. The fMRI signals have been scaled by the grand mean of each session. And we also compared the prediction accuracies of using ALFF and mALFF, which have yielded very few differences. The scaling may be effective for local voxels. But due to the spatial heterogeneity, the baseline variability may still present in some brain regions, which could be picked up by the machine learning algorithm. ReHo and connectivity measures use correlation measures, which are insensitive or have scaled local signal variability. This may explain why their prediction accuracies are smaller than ALFF. These problems may arise from the fact that most of the fMRI measures are relative measures. If absolute measures can be used, e.g. blood perfusion using arterial spin labeling (ASL) (Detre et al., 2012, 1992), then the effects from scanner stability may be minimized.

The current analysis demonstrated that machine learning is a powerful method that can pick up small effects. The phantom data showed the correlations between baseline MRI signals and daylight length were about 0.7. When scanning human participants, the background MRI signals outside the brain showed much smaller correlations with daylight length (Figure 5B and 5C). However, we could still achieve a similar level as the prediction accuracies by using machine learning as the phantom data (Figure 4). Indeed, the cross-validated prediction accuracies were between 0.6 and 0.7, which are very close to the correlation in the phantom data. Machine learning methods have become more and more popular in studying brain-behavior relationships (Cui and Gong, 2018; Finn et al., 2015) and brain alterations in mental disorders (Whelan et al., 2014). The current analysis illustrates that comparing performance with chance level may not be sufficient to control for potential confounding variables. Careful choice of control conditions is critical to make a proper conclusion. When performing machine learning analysis on functional activations or connectivity data, the structural MRI data may be a good choice as a control condition. The structural MRI data are usually available alongside the fMRI data, and don’t reflect the functional activity of the brain. Adding structural MRI as a control condition could rule out potential structural variations as a source of individual differences, but could also rule out potential MRI baseline variations as shown in the current analysis. A phantom scan may also be considered if long-term effects are of interest.

The current study did not completely rule out the potential seasonal or daylight effects on brain structures and functions. Studies using non-human animals have provided strong evidence of seasonal and daylight effects on brain structural and functional variation in hippocampal volume (Nissilä et al., 2012; Smulders et al., 1995; Tramontin and Brenowitz, 2000). PET studies of different neural transmitters also provide evidence of seasonal effects (Eisenberg et al., 2010; Kaasinen et al., 2012; Kalbitzer et al., 2010; Mc Mahon et al., 2016; Praschak-Rieder et al., 2008). Seasonal effects on brain functions may still exist, but it is difficult to study by using MRI due to the factors identified in the current analysis.

In conclusion, by applying machining learning on resting-state fMRI or structural MRI data, we can predict several environmental parameters, with the highest prediction accuracies to daylight length. However, the predictions were not likely due to the environmental effects on brain functions or structures, but may due to the baseline MRI signals. The data highlight the difficulty to use fMRI/MRI data to study long-term effects, and call for cautions to control for scanner stability when studying long-term effects.

## Acknowledgement

We thank Dr. James Hyde for his insightful comments on an earlier version of this manuscript. This analysis was funded by (US) National Institute of Health grants: R01AT009829 and R01DA038895.

