## Supplementary materials for "Estimations of the weather effects on brain functions using functional MRI: a cautionary note"

This PDF file include:

Figure S1 to S2

**S1. Effects of resting-state fMRI preprocessing**

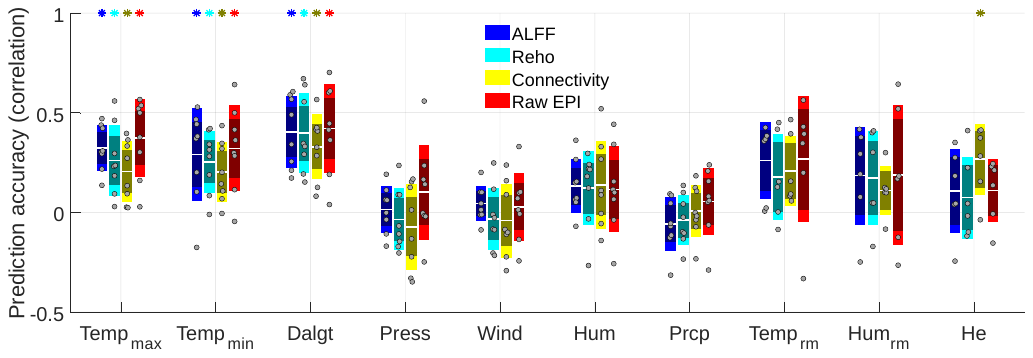

**Figure S1** Prediction accuracies (correlations) of the weather parameters using resting-state parameters with reduced preprocessing. ALFF refers to amplitude of low-frequency fluctuations; ReHo refers to regional homogeneity; and EPI refers to echo-planar imaging. Each dot represents one subject’s mean prediction accuracy. The center white lines, inner dark bars, and outer light bars represent the mean, 95% confidence interval, and standard deviation, respectively. The asterisks on the top represent statistical significance at p < 0.05 after false discovery rate (FDR) correction for all the 40 predictions.

**S2. Daylight length and MRI signals in the MPRAGE data**

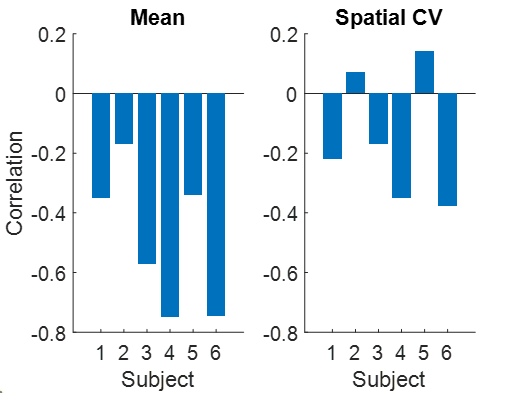

**Figure S2** The correlations between daylight lengths and MRI signals in the air mask in the six subjects from the Day2day dataset. Left, daylight length correlations with the mean MRI signals in the air mask. Right, daylight length correlations with the spatial coefficient of variation of the MRI signals in the air mask (standard deviation / mean).
